# Ecology dictates the value of memory for foraging bees

**DOI:** 10.1101/2021.09.06.458851

**Authors:** Christopher D. Pull, Irina Petkova, Cecylia Watrobska, Grégoire Pasquier, Marta Perez Fernandez, Ellouise Leadbeater

## Abstract

“Ecological intelligence” hypotheses posit that animal learning and memory evolves to meet the demands posed by foraging, and together with social intelligence and cognitive buffer hypotheses, provide a key framework for understanding cognitive evolution. However, identifying the critical environments where cognitive investment reaps significant benefits has proved challenging. Here, we capitalise upon seasonal variation in forage availability for a social insect model (*Bombus terrestris*) to establish how the benefits of short-term memory vary with resource availability. Through analysis of over 1700 foraging trips carried out over two years, we show that short-term memory predicts foraging efficiency – a key determinant of colony fitness – in plentiful spring foraging conditions, but that this relationship is reversed during the summer floral dearth. Our results suggest that selection for enhanced cognitive abilities is unlikely to be limited to harsh environments where food is hard to find or extract, highlighting instead that the complexity of rich and plentiful environments could be a broad driver in the evolution of certain cognitive traits.

## Introduction

The spatiotemporal distribution of food has been repeatedly theorized to contribute to the evolution of cognitive traits (1–5). In particular, the gross benefits of investment into learning and memory have been proposed to outweigh the significant constitutive and induced costs that these traits carry (6, 7) when food is hard to find because it is scarce, heterogenous, novel or challenging to extract (4, 8–10). Accordingly, comparative neuroanatomical studies have linked potentially demanding foraging tasks, such as remembering the location of cached food during high elevation winters, to changes in size or structure of neural regions across species (8, 11–13). However, the considerable challenges of standardizing confounding noncognitive factors (e.g. previous experience, parasite load or motivation) that can influence cognitive assay performance (14–16), when working with wild animals, mean that direct evidence to link cognitive abilities to fitness proxies is still rare. Those studies that have overcome such hurdles (e.g. 9, 14, 15) have not been extended to include variation across environments. As such, we do not yet have a full picture of when cognitive abilities are most valuable in the wild, and thus of the ecological conditions that favour cognitive evolution.

Here, we capitalize upon temporal variation in food availability within the colony lifespan of a social insect model to examine how the benefits afforded by a specific cognitive trait may vary with resource availability. *Bombus* are a temperate group and in most species workers begin to emerge in the early spring and colony foraging continues until late summer (19). The availability of floral forage varies considerably across this time, typically reaching a peak in the spring that recedes to a trough in late summer, with local variations (20). Foraging bees can visit hundreds of individual flowers across repeated foraging bouts every day, and cognitive traits such as learning and memory are considered fundamental in maximizing colony foraging success through their impacts on foraging efficiency (21). In laboratory settings, abundant evidence has documented the role of medium and long-term memory formation in the development of preferences for particular flower types or patches (22). Other work has shown that short-term memory (STM), a physiologically distinct memory phase that requires neither transcription nor translation and forms after a single trial, contributes to within-patch decisions (23–25). However, attempts to link memory to real-world foraging efficiency have focused on short summer foraging windows (18, 26), and so do not capture any variation in resource availability. In this study, we investigate the importance of STM for bumblebee foraging success across a highly variable foraging season.

Over two successive years, we reared 25 young, commercially supplied colonies (mean initial workers ± SD = 39 ± 16.8) under identical laboratory conditions following a staggered design from April to September (Fig. 1a). For each colony, a mean of nine (range 7-13) workers of known age underwent cognitive testing in a four-arm Radial Arm Maze (RAM; Fig. 1b). Originally developed for rodent toxicology, the RAM is a win-shift paradigm where performance is assayed as the number of revisits made to arms previously depleted of food within a foraging bout (for validation see Fig. S1). Bees were trained to asymptotic performance prior to testing (Fig. S2), such that cognitive assays required two weeks per colony. After cognitive testing was complete, tested workers were tagged with an RFID chip and screened for gut parasites – which may compromise cognition (27) – but no infections were found. Each colony was then given through-the-wall access to the external environment (Fig. 1c), comprising broadleaved mixed woodland and parkland surrounded by suburban housing and gardens (Fig. 1d). We recorded nectar and pollen foraging efficiency by individually weighing bees on entry and exit, until death, sampling ~ 47% of their total foraging career (*n* = 1735 nectar and pollen foraging trips). Of the 230 bees tested, 144 (63%) foraged in the wild.

**Figure 1.**
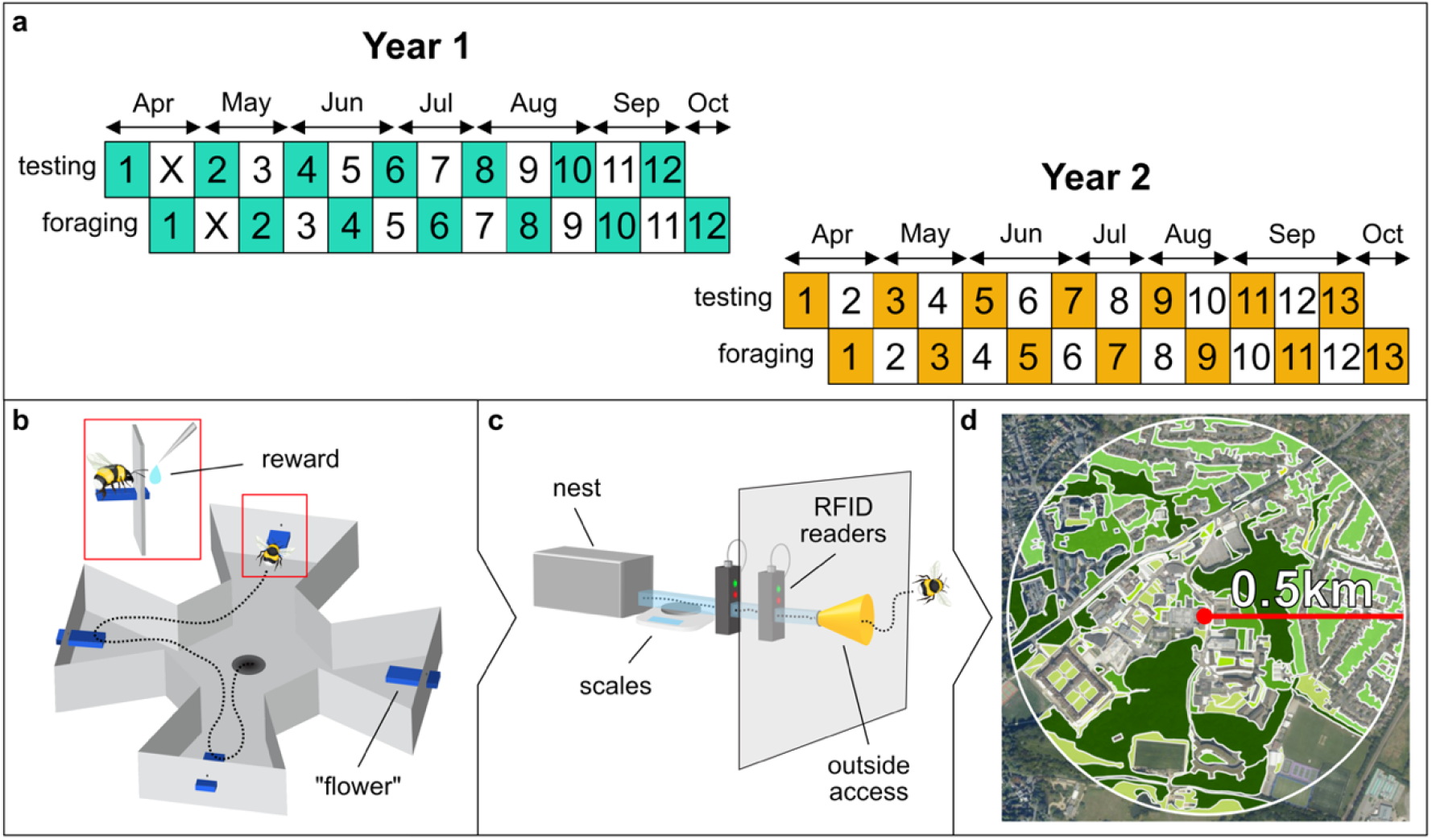
Lab-to-field testing of bee cognition and foraging efficiency. (*A*) Experimental timeline showing staggered design: each number indicates a colony (*n* = 25; colony X died during testing). (*B*) Four-arm radial arm maze (RAM) used to assay short-term memory (STM); inset shows bee extracting a reward. (*C*) Bees were weighed on exit and return to the nest, passing through RFID readers. (*D*) Colonies were surrounded by university parkland, suburban greenspace, and private gardens. The most dominant land type was mixed woodland (dark green). Non-coloured areas represent areas unlikely to contain floral resources.

## Results

We found that for nectar-collecting trips, the relationship between STM and foraging efficiency reversed in direction across the foraging season (Fig. 2 and Table S1; linear mixed effects model [LMER] containing RAM score x week interaction: ΔAIC to next best model = 4.7; interaction estimate ± 95% confidence interval = 0.05 ± 0.02 to 0.09). However, contrary to our expectation that STM would be most beneficial when resources were sparse, bees with better STM collected more nectar/minute in spring but less in summer, compared to bees with poorer RAM scores. Foraging efficiency also varied within individual lifetimes, whereby bees exhibited an increase in nectar foraging efficiency at the start, and a decrease towards the end, of their foraging careers, presumably as they gained experience and then underwent senescence, in accordance with previous results (19) (Figure S3a; estimate ± 95% CI = –0.03 ± –0.04 to –0.02). Larger bees were also more efficient foragers (Figure S3b; estimate ± 95% CI = 0.34 ± 0.15 to 0.52) and all bees foraged more efficiently on cooler, more humid days (Fig. S3c; estimate ± 95% CI = 0.01 ± 0.01 to 0.02). Neither year of the experiment (Fig. S3d; estimate ± 95% CI = –0.28 ± –0.58 to 0.04) nor bee age at initial release (Fig. S3e; estimate ± 95% CI = –0.03 ± −0.06 to 0.001) had significant impacts. Pollen foraging efficiency is not expected to vary as drastically with STM as nectar foraging, because flowers typically contain more pollen than can be extracted in a single visit (28). Avoidance of visited flowers within patches is thus less relevant and, accordingly, STM did not predict pollen foraging efficiency (Table S2; generalized linear mixed effects model [GLMER]: ΔAIC between intercept-only null model and next best alternative = 2.66).

**Figure 2.**
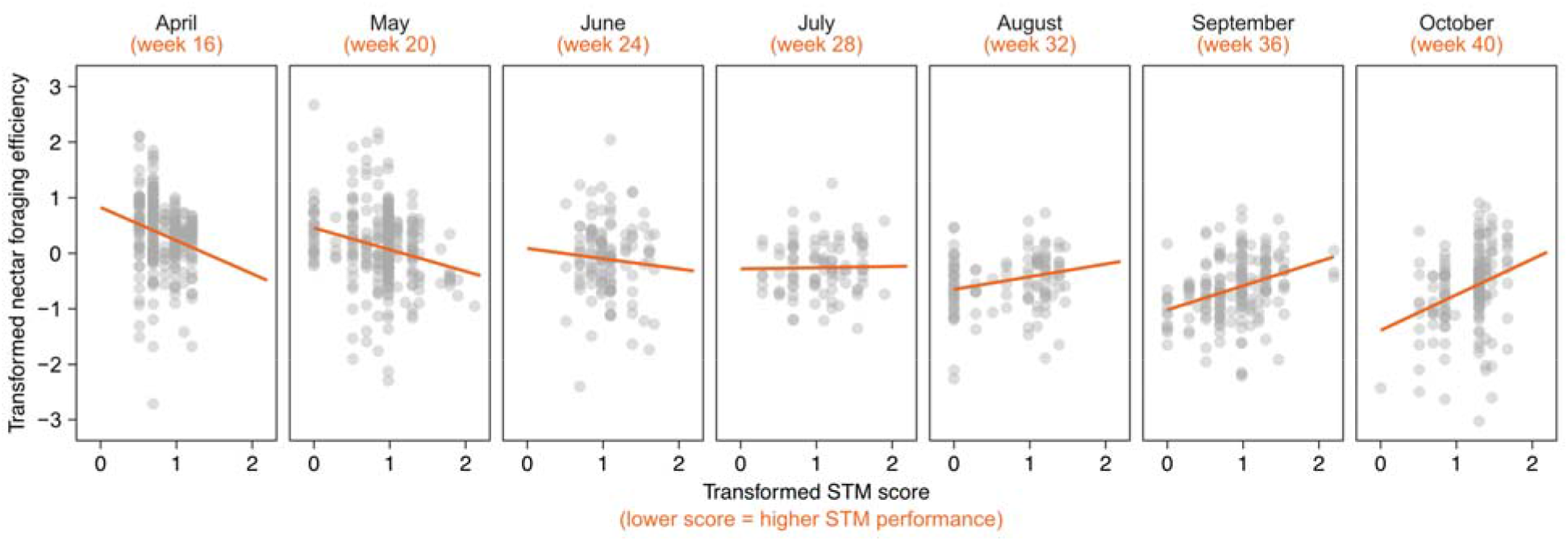
Seasonal reversal in bee STM and foraging efficiency relationship. Partial residual plots from a linear mixed effect model including an interaction between bee STM score and week of year (*n* = 1202 nectar foraging trips by 134 bees). Fitted lines indicate the relationship between STM and nectar foraging efficiency at four-week intervals, holding the effect of other numeric predictors constant at their median (covariates) or mode (factors). Both variables are presented, as analysed, on transformed scales (ORQ normalization and log(n+1), respectively). Negative nectar values result from bees leaving with more nectar than they return with (19).

Since our bees were RFID-tagged, we could also relate RAM performance to survival and total lifetime foraging effort (total number of trips), again across the whole season. However, both variables were solely predicted by age, with bees that were older on release dying sooner (cox proportional hazard model: estimate ± 95% CI = 0.07 ± 0.01 to 0.12) and thus conducting fewer trips (GLMER: estimate ± 95% CI = –0.35 ± –0.55 to –0.15). Adding RAM score to the models did not sufficiently improve fit in either case (Tables S3-4). Additionally, comparisons of tested and non-tested control bees revealed that cognition testing itself had no measurable impact on either the foraging performance or survival of bees (Tables S5-7).

Our results suggest that, rather than enhancing foraging efficiency in summer conditions, the benefits of STM were realised in spring. It is unlikely that this finding is the result of differences arising from confounding variables typically thought to influence cognition studies (14–16, 29): all colonies were raised from laboratory-raised queens in enclosed facilities under identical aseasonal conditions, bees were free from gut parasites during cognitive testing, there were no differences in prior experience, bees were of known age and size, and hunger-or condition-driven motivational effects were effectively standardized because workers forage to meet the demands of the colony. Instead, we believe that our findings are due to shifting patterns of floral resources, which are typically abundant in spring but limited in summer (20). To confirm this expected seasonal pattern in phenology, we combined land classification techniques with transect and quadrat sampling to measure floral generic richness within a 500m radius around our colonies, spanning mixed broadleaved woodland and parkland, local public green spaces, and residential gardens. Weekly surveys revealed an overall decline in the number of genera in flower towards the end of the season for all land types except grassland, which did not change (Fig. 3a; all statistics given in Table S8 and 9). Floral resources in wooded areas, which is the dominant land type in our survey area (woodland = 48%, open woodland = 11%), showed the sharpest linear decline from a peak in spring; others, such as private gardens (26%), increased towards a peak in early summer before declining by the end of summer. This pattern is also mirrored in the gross foraging trends found for all bees across the season: nectar and pollen foraging efficiency peaked in spring and was lowest at the end of summer, albeit with a brief revival in early autumn (Fig. 3b-c). Bout duration (Fig. 3d), which correlates with resource availability in bumblebees (30), followed the same pattern. Finally, identification of pollen loads collected by bees in the second year of our experiment showed a gradual decline in floral generic richness from a spring peak of fifteen genera to a late summer trough of one genus (Fig. S4). Together, these data indicate that spring foraging environments were plentiful in comparison to the relatively depauperate summer.

**Figure 3.**
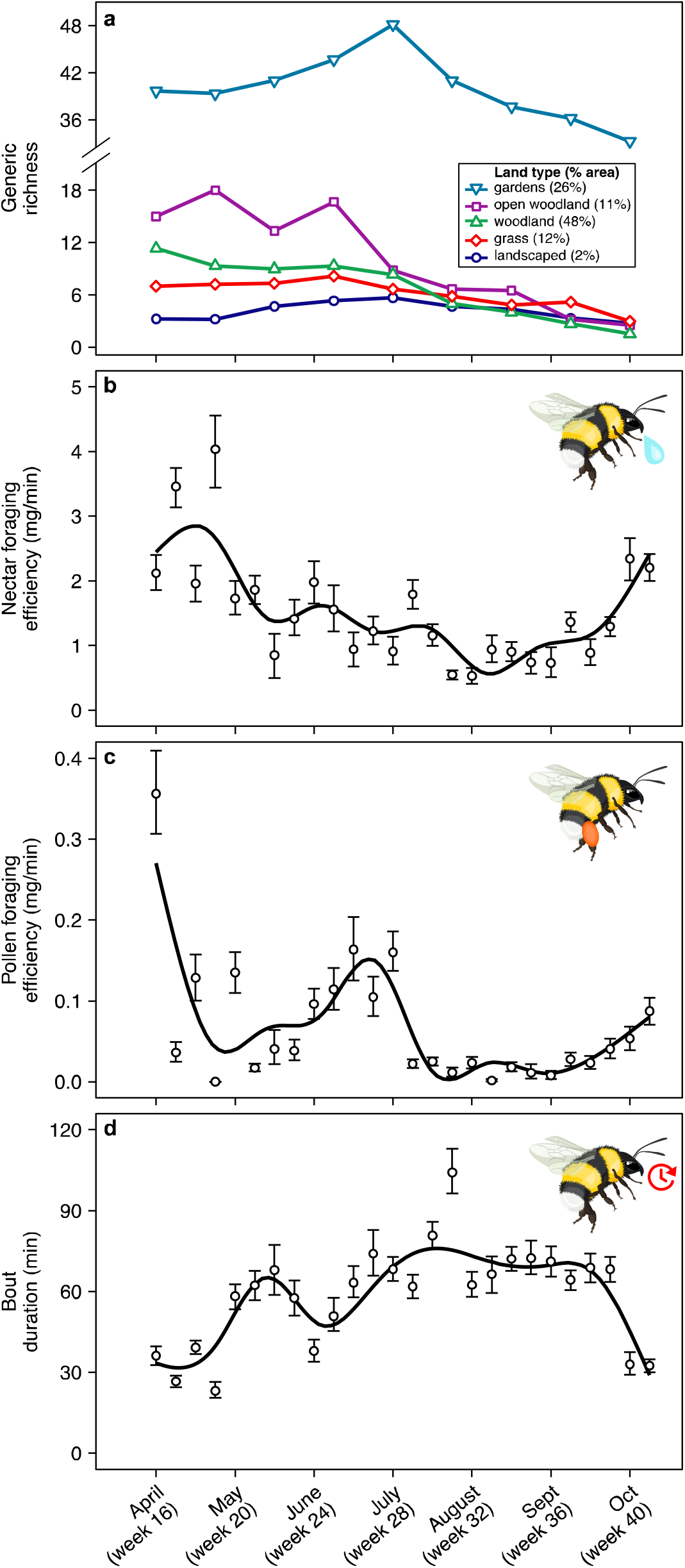
Seasonal variation in floral resources and foraging. (*A*) Shifts in floral generic richness across the foraging season. Each point represents a three-week average (or two weeks for the final survey) based on both years of the study (except surveys 1-4 for *gardens, open woodland* and *woodland*; see methods). Percentages in legend indicate proportion of each land type within the survey area. (*B-D*), Nectar foraging efficiency, pollen foraging efficiency and bout duration for whole colonies (*n* = 6616 foraging trips) across both years of the foraging season (weekly mean ± 95% CI; trend lines fitted using a generalized additive model (GAM) smoother function).

## Discussion

Our finding that RAM performance was associated with higher foraging efficiency in plentiful spring conditions rather than harsh summers is surprising, given that food scarcity (e.g. high-elevation winters (9, 31)) is more typically associated with cognitively demanding foraging. Learning about and remembering the floral characteristics that predict reward would seem likely to reap more benefits in sparse conditions when fewer floral resource patches are available, and travel times between them are higher. However, this result is nevertheless consistent with the nature of the within- (*c.f*. between-) patch foraging task that the RAM seeks to mirror. For social bees, foraging is characterised by an initial trip from the nest to the first flower patch, and then short flights from flower to flower within a patch (such as a flowering tree or shrub) followed, once rewards diminish, by longer between-patch trips before returning to the colony. The ability to remember, and thus avoid, recent visits within a patch is thought to contribute significantly to within-patch efficiency (19), while between-patch efficiency is more likely to depend upon longterm memories of rewarding locations. In spring, when resource patches are rich and plentiful near to the colony, within-patch efficiency effects may have greater impact than in summer, when reaching and moving between floral patches requires long travel times that dwarf any within-patch savings. Accordingly, previous work in the same species (18) has shown that performance in a task requiring longer-term memory predicted foraging efficiency at the height of summer (but see (26)).

A decrease in the relative importance of within-patch efficiency savings with patch density could explain why high-scoring bees performed relatively well in spring, but not why STM was detrimental (rather than simply inconsequential) in summer. Why should strong STM performance come at a cost to foraging efficiency, rather than simply producing benefits that are less detectable, when resources are sparse? One possibility is that investment in STM comes at a cost to other cognitive traits. In *Drosophila*, the short-term ability to form single-trial memories (or multiple trials with very short inter-trial intervals – termed anaesthesia-resistant memory ARM) – trades off against the long-term memory (LTM), such that flies with good ARM have poorer LTM (32). However, this trade-off has not yet been explored in bees. Alternatively, STM investment may trade-off against physiological or metabolic traits, given that investment into cognition has been found to come at significant constitutive and induced costs (7, 33). We found no significant effect of STM on individual survival, but we cannot rule out the possibility of sublethal costs of cognitive investment that emerge in summer when flight times are relatively long, manifested through flight efficiency or the need to consume more of the nectar that is collected whilst foraging. Future research could productively investigate the link between such metabolic costs and investment in memory, because sublethal stressors can critically compromise colony reproductive success (34).

Although variation in the benefits afforded by cognitive traits have been hypothesized to vary with environmental conditions (1–4) – even within individual lifetimes – direct demonstrations of such fluctuations in potential selective value have not previously been performed. Here, we have provided evidence that the benefit of a cognitive trait can indeed vary considerably, even within a relatively short single foraging season, becoming apparent in plentiful, rich floral environments but reversing in the depauperate ones. For bumblebees, the timescale of this reversal did not fall within the expected lifespan of individual workers, who typically live for only a few weeks in the wild (19), but instead within the lifespan of the colony. Thus, future studies could evaluate the possibility that worker cognitive abilities may even vary with the stage in the foraging season at which an individual emerges, allowing bees to capitalize upon spring-flowering trees and hedgerows that are known to be key to colony survival and performance (35). In longer-lived animals that will experience variability within a single lifetime, such patterns might be expected to compromise selection for certain cognitive traits if the costs of maintenance in some periods outweigh the benefits in others.

While previous work within the ecological intelligence framework has placed emphasis on the potential role of food scarcity in driving cognitive evolution (5, 6, 9, 31), our study suggests that rich, plentiful environments, where food is easy to find, could be just as important a selective environment for certain cognitive traits. In our study, STM allowed individuals that could function especially efficiently within such complexity to excel at foraging, when all individuals fared relatively well. However, such effects will be specific to the cognitive trait in question, and LTM may well show the opposite pattern, both in our *Bombus* system and more generally. For example, in populations of black-capped chickadees (*Poecile gambeli*) that live at higher elevation and thus experience harsher winters, performance on a task that requires longer-term memory than the one-trial task described here correlates with overwinter survival (9). Likewise, a previous study in bumblebees that focused on summer conditions found that longer-term memory correlated with colony-level foraging efficiency (18). In both these cases, food is likely to be sparse and longer-term memories of cache/forage locations may well be more important than the short-term one-trial “working” memories that could improve within-patch efficiency. Overall, we have found strong support for the main tenet of the ecological intelligence framework: that the benefits of cognitive investment vary with the foraging environment. Our study thus points to the major role that ecological differences possibly play in generating and maintaining the considerable intraspecific cognitive variation observed across the animal kingdom (14, 15, 29).

## Materials and Methods

### Experimental overview

From April to October in 2018 and 2019, following a staggered design (Fig. 1a), we performed laboratory-based cognitive testing on 230 worker bees from 25 commercially sourced colonies (Fig. 1b; *n* = 12 in 2018 and 13 in 2019; one colony removed in April 2018 due to colony death pre-release). After a two-week testing period, tested bees were screened for gut parasites and RFID-tagged, and colonies were connected to a through-the-wall external access hatch (Fig. 1c). Nectar and pollen foraging efficiency was monitored for two weeks (approx. 6h per day, 4-5 days per week). At the same time, testing began for the next colony in the cycle (Fig. 1a). After foraging efficiency recording had ceased, colonies were permitted to continue foraging for another two weeks, during which time survival and foraging activity (but not foraging efficiency) was recorded through automated RFID readers. Throughout, we conducted weekly surveys of the land surrounding the foraging colonies, to record changes in the abundance and diversity of floral resources over the seasons (Fig. 1d).

### Bee colonies

We used commercial colonies of UK-native *Bombus terrestris audax* (BioBest, Waterloo, Belgium), housed in two-chamber plastic nestboxes (28 [l] x 16 [w] x10.5 [h] cm). Commercially supplied colonies are reared from domestic *Bombus* lines and queens had emerged as gynes, hibernated, and founded colonies in identical controlled indoor conditions. All colonies were young on arrival (mean initial number of workers ± SD = 39 ± 16.8), and only bees that emerged post-arrival (identified through tagging all bees with numbered discs on arrival and new bees after emergence [Abelo, UK]) were tested in our cognitive assay. Prior to cognitive testing, colonies were fed an *ad libitum* supply of inverted sugar syrup via an in-nest feeder (45% [w/v]; Thorne, Windsor, U.K.), with one 1.5g pollen ball (2:1 honeybee-collected pollen: 45% inverted sugar) added daily (two on Fridays; none on weekends). During cognitive testing, pollen feeding continued, but syrup feeders were removed and between 10-15 ml (depending on colony size and stores) was pipetted into nectar pots per weekday evening. Colonies were not fed once given outside access.

### Radial arm maze

We used a radial arm maze (Fig. 1b) to assess the STM of individual bees. The RAM is a win-shift paradigm in which all arms are initially baited with food rewards that are not replaced upon removal, within one bout. Revisits to depleted arms constitute “errors” and the number of errors within a single foraging bout is the measure of STM (36). The RAM is therefore an ecologically relevant task that mimics natural nectar foraging within a flower patch. Our RAM consisted of an octagonal four-arm array in which differentiation of the arms was possible through a laminated black and white panoramic image of the laboratory at the ends of each arm. Rewards were accessed via removable blue, rectangular Perspex platforms – henceforth “flowers” – at the end of each arm (colour number = 744; 7.5 [l] x 3 [w] x 0.5 [h] cm) that slotted through holes in the maze wall (Fig. 1b). Bees retrieved a sucrose reward (see “Cognitive testing”) by alighting on flowers and inserting their proboscises through small holes in the RAM wall (Fig. 1b inset).

### Cognitive testing

Cognitive testing began 12 days after arrival in the laboratory (to allow bees of known age to emerge) and commenced with a group training period (~1h) whereby bees were allowed to forage freely in the RAM and all arms were continually rewarding (an *ad libitum* supply of 2M sucrose solution), and to enter and leave the arena at will. Motivated foragers were then tested in the RAM alone for 12 foraging bouts, during which all arms were baited with 20 μl of 2M sucrose solution that was not replaced within a bout (except the last non-depleted flower when bees were fed to repletion). After each landing, flowers were replaced with identical clean, unrewarded replacements to preclude the use of scent marks. Access to the nest during testing was prevented via sliding shutters, unless a bee spent >30 s trying to return or attempted to return more than twice within a bout (to maintain foraging motivation). Consequently, in some bouts not all arms were visited. All bouts were filmed for later video analysis using BORIS video analysis software (37).

### Parasite screening and RFID chips

We screened faecal samples from all tested bees for the presence of gut parasites (*Apicystis bombi, Nosema bombi* and *Crithidia bombi*) at the end of the two weeks of cognition testing (Nikon e50i) and measured intra-tegula distance as a proxy for overall size, as in other studies (e.g. (18, 25)). An RFID chip (Microsensys GmbH) encoding a unique 16-digit identification number was also superglued (Loctite) to the thorax. We performed the same procedure for a cohort of non-tested bees from each colony that were observed foraging during group training (mean of four bees per colony, total *n* = 99), to confirm that our testing regime had no effect on subsequent foraging efficiency.

### Measuring foraging performance

We measured the foraging success of individual bees on exiting and re-entering the colony using weight-averaging scales for moving subjects (mean of three repeat measurements with 2s averaging and accuracy of ± 2 mg; Advanced portable balance Scout STX123 120g; OHAUS Corporation) and their lifetime foraging activity and survival using an RFID system (MicroSensys GmBH, Fig. 1c). Per trip nectar values were calculated by subtracting the bee’s outbound weight from her inbound weight, minus the weight of any pollen. We removed pollen from one leg of the bee – via a trap door in the tube – and weighed and froze (−20) the pollen for later identification (see below). The weight of the pollen was doubled and subtracted from the bees’ weight. On entry and exit, bees passed through two RFID readers (Microsensys GmbH) that recorded identity and travel direction. We collected foraging performance data for ~ 6 hours per day, four-five days a week, for two weeks per colony. In total, 6616 were trips recorded across all colonies and bees. For analysis, nectar trips (*n* = 1202 trips by 134 bees) were defined as those where < 3 mg of pollen was collected, and pollen trips > 3 mg (*n* = 526 trips by 91 bees), based on histograms of foraging data. Any trips that were less than 7 min in duration were not included in the analysis, because such trips are more likely to represent orientation flights or waste disposal.

Colonies then foraged for a further two weeks with only RFID data collection to assay survival, in which time >99% of RFID-chipped bees eventually failed to return to the colony and were presumed dead. We affixed brightly coloured plastic cones onto the outdoor entrance of nests and replaced pollen that we removed daily for microscopic pollen identification using a local reference collection with equal quantities of honeybee collected pollen (Koppert, UK). Colonies were euthanized at the end of four weeks of foraging.

### Pollen identification

Pollen loads were defrosted, vortexed for 60 s and suspended in water (1 mg pollen:10 μl pure water); 1 μl was added to a glass slide and heated at 50 °C for 20 s, followed by two drops of melted glycerine with fuschine dye (Brunel Microscopes). The slide was covered with a slip and left to cure at 50 °C for 30 seconds. We discounted any floral morphotype where the grain count was < 50 grains per section of slide that was counted. Morphotypes were microscopically identified to genus level using a combination of sources (38, 39) and the pollen reference collection at Royal Holloway University of London.

### Floral resource surveys and weather data

We used QGIS to (*i*) classify the 500 m radius surrounding our colonies into broad land use types likely to contain forage: woodland, grassy woodland, grassland and landscaped and (*ii*) select sampling sites within these categories. Additionally, we arranged access to private suburban gardens (total sites per land use type = 12, except grass = 24 and gardens = 11). Surveys were performed for one day each week, whereby each site was visited once every three weeks on a rotational basis. Methods were customized to each land-use type: in grass and landscaped areas, we sampled 0.25m^2^ quadrats; in woodland and open woodland we surveyed 30 m transects; in gardens we counted every genus of plant in flower. In year one, we utilized a different approach to record transect and garden data that was discarded after the 4^th^ survey; hence we only include transect and garden data from year two in survey blocks 1-4 of Fig. 3a. A three-week average was calculated using generic richness data from all three sets of sites for the same survey period in both years for graphical representation in Fig. 3a. Local meteorological data was collected from a weather station located ~ 8 km away at Imperial College London, Silwood Park. Hourly weather data (temperature, humidity, and wind speed) was averaged to produce a daily mean for analysis.

### RAM validation

In an additional experiment, we tested whether bees perform better on the RAM than using stereotypical movement rules or by chance alone. Using the same procedure as before, we tested 20 bees from four colonies on an eight-arm version of our RAM (to increase the difficulty of the task). Each bee performed ten training bouts, to learn how to use the maze, followed by ten test bouts once performance had plateaued. Following (23, 25), we used data from the test bouts to calculate the general probability of moving from each flower to each of the other flowers (e.g. from F1 to F2, from F1 to F3 etc.), creating an individual transition matrix for each bee. Following a Monte Carlo process, we then simulated each bee’s expected performance following solely this matrix, and not avoiding visited flowers (10000 iterations per bee). A similar process was followed to estimate likely performance following random selection of platforms.

### Statistical analysis

All statistical analyses were conducted using R version 4.1.0 and an information-theoretic approach. For foraging and survival models, we built a candidate model that contained every hypothesized covariate (see below and Supplementary Tables 1-10 for model descriptions) and compared this to (*i*) the same model, but with STM score as a predictor, (*ii*) the same model but including an interaction between STM score and week (*iii*) a null model containing only the intercept and random factors. Model selection was based on ΔAIC or ΔAICc (depending on sample size), where a cut-off of >2 was used to identify the best model. If nested models had comparable fit the simplest model was selected.

We modelled nectar foraging efficiency using a linear mixed effects regression (LMER) that included foraging efficiency as the response (mg/min; ordered quantile normalization transformation using the BestNormalize package (40) due to heteroscedasticity) and week-of-year, STM score (log(n+1) transformed to reduce influence of outliers) and their interaction as our predictors of interest, with initial bee age (at time of release), bee size, year of experiment, foraging experience (number of days since foraging began), and a composite “weather” score as covariates. The composite weather score was produced via principal component analysis to reduce temperature, humidity, and wind speed into a single component (~ 84% variation captured). To account for collinearity between weather scores and week, weather scores in the main analysis are the residuals of generalized additive model (GAM) predicting weather based on week; thus, any reported effects of “weather” reflect those that occurred in addition to effects of week. We included foraging experience as a quadratic polynomial to account for its non-linear effect, which improved model fit. For each bee, we fit random intercepts that interacted with random slopes for experience (because improvement in performance could vary between bees and is likely affected by starting performance) and a random intercept per colony. To model pollen foraging efficiency, we used a generalised linear mixed effect regression (GLMER) with log-link Gamma errors; foraging experience as a linear predictor and uncorrelated random slopes and intercepts fit this data best, otherwise the model was identical to the nectar LMER.

To analyses survival data, we used a Cox proportional hazards model with week of year, STM score and their interaction as our predictors of interest, and bee age (at time of release), bee size, composite weather score, and year of experiment as covariates, including colony as a random effect using the shared frailty function. We used a GLMER with Poisson error distribution to model lifetime foraging effort, including number of RFID-recorded trips as the response and an observation-level random effect to account for overdispersion (41); all other covariates were identical to the nectar foraging model and colony was included as a random intercept.

To analyse generic floral richness, we built linear regressions that included weekly generic richness as the response, season as a linear predictor or quadratic polynomial (depending on fit), and year of experiment as a covariate. For gardens and woodland data, we applied a log(n) and ORQ transformation respectively to the response to achieve normality in model residuals.

We analysed the RAM performance of bees (*n* = 230) using a GLMER with negative binomial error distribution (due to overdispersion). We included the per bout number of errors (revisits to previously depleted flowers) as the response and bout number (1–12) as the predictor of interest, with bee size, the age of the bee, and whether the bee had engaged in group training on the morning of testing (see main methods) as covariables. We allowed the bees’ intercepts, and the slope for bout, to vary individually, whilst also including a correlation between the intercept and slope. Additionally, we included colony of origin as a random.

We built several models to investigate if our cognition testing regime had a subsequent impact on the foraging performance and survival of tested bees. Firstly, we fit a LMER with Gaussian error distribution that had nectar foraging efficiency (mg/min) per foraging trip as the response, with treatment (control or cognition tested) as a main effect, and bee size, initial age, year of experiment, composite weather score, days since release as a quadratic polynomial and week of year as covariates. We transformed the response using QRQ normalization due to heterogeneity. We fitted random intercepts per bee that interacted with random slopes for experience and a random intercept per colony. Secondly, we ran a GLMER with Gamma error distribution and log link that had pollen foraging efficiency (mg/min) per foraging trip as the response, and the same covariates as above, except day since release was included as a linear term and the interaction between the random intercepts and slopes was removed. Lastly, we fit a cox proportional hazards model that included treatment as a main effect and bee size, initial age, composite weather score, year of experiment and week as covariates, with colony as a random effect using the frailty function.

For mixed effects modelling, we used the ‘lme4’ package (42) and checked all model assumptions via residual plots and functions in ‘influence.ME’ and ‘performance’ packages (43, 44). For the survival model, we plotted residuals, tested for non-proportional hazards, and assessed the model for influential observations.

## Acknowledgments

We are grateful to Rosie Wright, Matthew Rawlings, Monika Yordanova, Harry Siviter, Matthew Hasenjager and Fabio Manfredini for help with data collection, the Egham and Englefield Green residents who provided access to their gardens for floral surveys, and Imperial College London, Silwood Park for providing access to weather station data. This research was supported by a Leverhulme Trust Research Project Grant to E.L. (RPG-2016-444) and C.W. was funded through the IC-RHUL BBSRC DTP (BB/M011178/1).

## Notes

### Competing Interest Statement

The authors have declared no competing interest.

