## Supplementary Information for "Ecology dictates the value of memory for foraging bees"

Christopher D. Pull.

**Fig. S1.** Bees perform better than chance alone or by using stereotypical movement rules on an eight-arm radial arm maze. For simulated estimates, vertical lines represent mean values whilst the red vertical line represents the intercept of a GLMM with Poisson error structure, colony as a fixed effect, and a random intercept per bee for the observed data; frequency bars represent the intercepts from the same model obtained from the 10000 simulated datasets (*n* = 20 bees).

**Fig. S2.** Bee performance on a four-arm RAM. Number of errors during the last three test bouts (10-12) were averaged to produce a mean STM score for subsequent analysis. Dots ± error bars show mean number of errors per bout ± 95% CIs (n = 230 bees). Bees showed improved RAM performance with increasing bout number, indicating learning (Supplementary Table 10; ∆AIC between full and next best model = 76.46; effect of bout [estimate ± 95% confidence internal] = –0.07 ± –0.08 to ­–0.05). Group training on the morning of testing also improved STM performance (estimate ± 95% CI = 0.14 ± 0.02 to 0.25), whereas size (estimate ± 95% CI = 0.06 ± –0.03 to 0.15) and age (estimate ± 95% CI = 0.003 ± –0.01 to 0.02) had no impact.

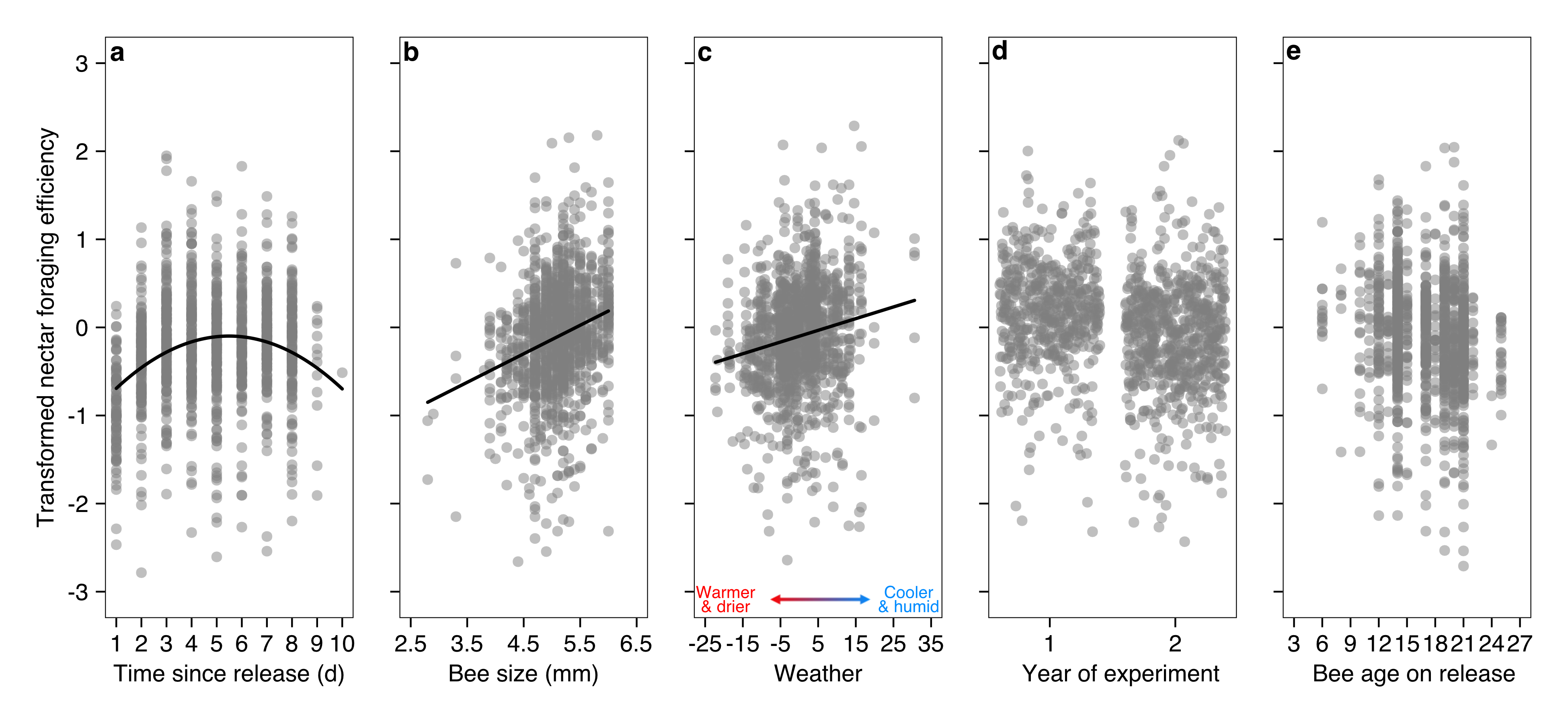

Fig. S3. Influence of covariates on nectar foraging efficiency. Partial residual plots for covariates from a linear mixed effect model with an interaction between bee STM score and week of year (*n* = 1209 nectar foraging trips; STM x week interaction displayed in Fig 2). Fitted lines (a-c) included where there is significant relationship between the covariate and nectar foraging efficiency, while holding the effect of other numeric predictors constant at their median and by setting year to “two” (most common value). Nectar foraging efficiency is presented, as analyzed, on transformed scales (ORQ normalization); for reference, untransformed nectar values range from -6.75–14.85 mg/min.

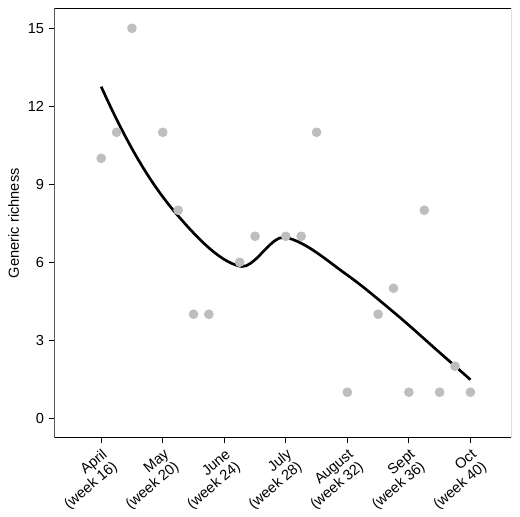

Fig. S4. Generic richness of pollen samples collected from cognition-tested bees in second year of experiment. Dots represent summed generic richness values for the week pollen samples were collected in, and smoothed trend line was fitted via LOESS.

Table S1. Nested candidate model set and model selection outcome for an LMER investigating the relationship between season (week of year) and STM score on nectar foraging efficiency. All models included the same random effect structure and, except the intercept-only null model, the same set of covariates. An ∆AIC > 2 was used to select the final model (highlighted in bold).

| **Covariates:** size + initial age + year + composite weather + quadratic polynomial (days since release)  **Random effect structure:** (days since release \| bee) + (1 \| colony) | | | | | |
| --- | --- | --- | --- | --- | --- |
| Model | DF | Log likelihood | AIC | ∆AIC | Akaike weight |
| **week * stm** | **15** | **-1318.27** | **2666.53** | **0.00** | **0.89** |
| - stm | 13 | -1322.62 | 2671.23 | 4.70 | 0.08 |
| week + stm | 14 | -1322.61 | 2673.23 | 6.70 | 0.03 |
| null | 6 | -1359.31 | 2730.62 | 64.09 | 0.00 |

Table S2. Nested candidate model set and model selection outcome for a GLMER investigating the relationship between season (week of year) and STM score on pollen foraging efficiency. All models included the same random effect structure and, except the intercept only null model, the same set of covariates. An ∆AIC > 2 was used to select the final model (highlighted in bold).

| **Covariates:** size + initial age + year + composite weather score + days since release  **Random effect structure:** (1 \| bee) + (0 + day since release \| bee) + (1 \| colony) | | | | | |
| --- | --- | --- | --- | --- | --- |
| Model | DF | Log likelihood | AIC | ∆AIC | Akaike weight |
| **null** | **5** | **328.480** | **-646.96** | **0.00** | **0.70** |
| - stm | 11 | 333.149 | -644.30 | 2.66 | 0.19 |
| week + stm | 12 | 333.302 | -642.60 | 4.40 | 0.08 |
| week * stm | 13 | 333.538 | -641.08 | 5.88 | 0.04 |

**Table S3.** Nested candidate model set and model selection outcome for a Cox proportional hazards model investigating the relationship between season (week of year) and STM score on bee survival. All models included the same frailty function for non-independency and, except the intercept-only null model, the same set of covariates. As two models were within 2 ∆AIC points, the simplest (least terms) was chosen as the final model (highlighted in bold).

| **Covariates:** size + initial age + year + composite weather score  **Random effect structure:** frailty (colony) | | | | | |
| --- | --- | --- | --- | --- | --- |
| Model | DF | Log likelihood | AIC | ∆AIC | Akaike weight |
| **- stm** | **12** | **546.14** | **1116.96** | **0.00** | **0.51** |
| week * stm | 15 | 544.32 | 1118.85 | 1.89 | 0.20 |
| week + stm | 12 | 546.67 | 1119.07 | 2.11 | 0.18 |
| null | 10 | 548.94 | 1119.82 | 2.87 | 0.12 |

**Table S4.** Nested candidate model set and model selection outcome for a GLMER investigating the relationship between season (week of year) and STM score on lifetime foraging effort (total number of bouts recorded by RFID system). All models included the same random effect structure and, except the intercept only null model, the same set of covariates. As two models were within 2 ∆AIC points, the simplest (least terms) was chosen as the final model (highlighted in bold).

| **Covariates:** size + initial age + year  **Random effect structure:** (1 \| colony) | | | | | |
| --- | --- | --- | --- | --- | --- |
| Model | DF | Log likelihood | AIC | ∆AIC | Akaike weight |
| **- stm** | **7** | **-689.87** | **1394.58** | **0.00** | **0.52** |
| week * stm | 9 | -688.38 | 1396.13 | 1.55 | 0.24 |
| week + stm | 8 | -689.87 | 1396.82 | 2.24 | 0.17 |
| null | 3 | -696.11 | 1398.39 | 3.82 | 0.08 |

**Table S5.** Nested candidate model set and model selection outcome for a LMER investigating the effect of cognition testing on the nectar foraging efficiency of RAM tested and control bees. All models included the same random effect structure and, except the intercept only null model, the same set of covariates. As two models were within 2 ∆AIC points, the simplest (least terms) was chosen as the final model (highlighted in bold).

| **Covariates:** size + initial age + year + composite weather + quadratic polynomial (days since release) + week  **Random effect structure:** (days since release \| bee) + (1 \| colony) | | | | | |
| --- | --- | --- | --- | --- | --- |
| Model | DF | Log likelihood | AIC | ∆AIC | Akaike weight |
| **- treatment** | **13** | **-2222.39** | **4470.78** | **0.00** | **0.51** |
| + treatment | 14 | -2221.42 | 4470.84 | 0.059 | 0.49 |
| null | 6 | -2285.48 | 4582.96 | 112.18 | 0.00 |

**Table S6.** Nested candidate model set and model selection outcome for a LMER investigating the effect of cognition testing on the pollen foraging efficiency of RAM tested and control bees. All models included the same random effect structure and, except the intercept only null model, the same set of covariates. As two models were within 2 ∆AIC points, the simplest (least terms) was chosen as the final model (highlighted in bold).

| **Covariates:** size + initial age + year + composite weather + days since release + week  **Random effect structure:** (1 \| bee) + (0 + day since release \| bee) + (1 \| colony) | | | | | |
| --- | --- | --- | --- | --- | --- |
| Model | DF | Log likelihood | AIC | ∆AIC | Akaike weight |
| **null** | **5** | **499.29** | **-988.58** | **0.00** | **0.74** |
| - treatment | 11 | 466.92 | -985.83 | 2.76 | 0.19 |
| + treatment | 12 | 466.95 | -983.83 | 4.76 | 0.07 |

**Table S7.** Nested candidate model set and model selection outcome for a Cox proportional hazards model investigating the effect of cognition testing on the survival of RAM tested and control bees. All models included the same random effect structure and, except the intercept only null model, the same set of covariates (see methods). As two models were within 2 ∆AIC points, the simplest (least terms) was chosen as the final model (highlighted in bold).

| **Covariates:** size + initial age + year + composite weather score + week  **Random effect structure:** frailty (colony) | | | | | |
| --- | --- | --- | --- | --- | --- |
| Model | DF | Log likelihood | AIC | ∆AIC | Akaike weight |
| **- treatment** | **15** | **-1048.92** | **2130.52** | **0.00** | **0.65** |
| + treatment | 16 | -1048.44 | 2131.78 | 1.26 | 0.35 |
| null | 13 | -1057.13 | 2142.06 | 11.54 | 0.002 |

**Table S8.** Nested candidate model set and model selection outcome for linear regressions investigating the effect of season (week of year) and year of experiment on floral generic richness across survey land types. An ∆AIC > 2 was used to select the final model (highlighted in bold); if more than one model was within 2 ∆AIC points, then the simplest (least terms) was chosen.

| Land type | Model | DF | Log likelihood | AICc | ∆AIC | Akaike weight |
| --- | --- | --- | --- | --- | --- | --- |
| Gardens | **- year** | **4** | **0.991** | **7.20** | **0.00** | **0.56** |
|  | week + year | 5 | 1.405 | 9.00 | 1.79 | 0.23 |
|  | null | 2 | -2.431 | 9.20 | 2.02 | 0.21 |
| Grass | **week + year** | **5** | **-105.00** | **221.4** | **0.00** | **0.90** |
|  | - year | 4 | -108.62 | 226.4 | 4.75 | 0.08 |
|  | null | 2 | -112.27 | 228.8 | 7.41 | 0.02 |
| Landscaped | week + year | 5 | -97.74 | 206.9 | 0.00 | 0.58 |
|  | **- year** | **4** | **-99.39** | **207.7** | **0.81** | **0.39** |
|  | null | 2 | -104.12 | 212.5 | 5.62 | 0.04 |
| Open woodland | **week + year** | **4** | **-98.89** | **206.9** | **0.00** | **1** |
|  | - year | 4 | -109.22 | 227.6 | 20.64 | 0 |
|  | null | 2 | -127.74 | 259.8 | 52.88 | 0 |
| Woodland | **- year** | **3** | **-39.12** | **84.9** | **0.00** | **0.76** |
|  | week + year | 4 | -39.02 | 87.2 | 2.27 | 0.24 |
|  | null | 2 | -55.29 | 114.9 | 30.00 | 0.00 |

**Table S9.** Parameter estimates ± 95% CIs for the best fitting model per land type.

| Land type | Parameter | Estimate | Lower | Upper |
| --- | --- | --- | --- | --- |
| Gardens | quadratic polynomial (week) | -0.001 | -0.003 | -0.0002 |
| Grass | quadratic polynomial (week) | -0.01 | -0.03 | 0.00007 |
|  | year | 1.66 | 0.41 | 2.92 |
| Landscaped | quadratic polynomial (week) | -0.01 | -0.03 | -0.006 |
| Open woodland | season | -0.50 | -0.65 | -0.35 |
|  | year | 5.41 | 3.23 | 7.61 |
| Woodland | season | -0.10 | -0.13 | -0.07 |

**Table S10.** Nested candidate model set and model selection outcome for a GLMER investigating the relationship between STM errors and bout number whilst solving the RAM. All models included the same random effect structure and, except the intercept only null model, the same set of covariates (see methods). An ∆AIC > 2 was used to select the final model (highlighted in bold).

| **Covariates:** size + age tested + group training  **Random effect structure:** (bout number \| bee) + (1 \| colony) | | | | | |
| --- | --- | --- | --- | --- | --- |
| Model | DF | Log likelihood | AIC | ∆AIC | Akaike weight |
| **+ bout number** | **10** | **5274.69** | **10569.38** | **0.00** | **1** |
| - bout number | 9 | 5313.92 | 10645.83 | 76.46 | 0 |
| null | 6 | 5317.77 | 10647.54 | 78.16 | 0 |
